# A neurocomputational model for intrinsic reward

**DOI:** 10.1101/2019.12.19.882589

**Authors:** Benjamin Chew, Bastien Blain, Raymond J Dolan, Robb B Rutledge

**Affiliations:** Max Planck University College London Centre for Computational Psychiatry and Ageing Research, London WC1B 5EH, United Kingdom; Wellcome Centre for Human Neuroimaging, University College London, London WC1N 3BG, United Kingdom; Department of Psychology, Yale University, New Haven, CT 06520, USA

## Abstract

Standard economic indicators provide an incomplete picture of what we value both as individuals and as a society. Furthermore, canonical macroeconomic measures, such as GDP, do not account for non-market activities (e.g., cooking, childcare) that nevertheless impact well-being. Here, we introduce a computational tool that measures the affective value of experiences (e.g., playing a musical instrument without errors). We go on to validate this tool with neural data, using fMRI to measure neural activity in male and female human subjects performing a reinforcement learning task that incorporated periodic ratings of subjective affective state. Learning performance determined level of payment (i.e., extrinsic reward). Crucially, the task also incorporated a skilled performance component (i.e., intrinsic reward) which did not influence payment. Both extrinsic and intrinsic rewards influenced affective dynamics, and their relative influence could be captured in our computational model. Individuals for whom intrinsic rewards had a greater influence on affective state than extrinsic rewards had greater ventromedial prefrontal cortex (vmPFC) activity for intrinsic than extrinsic rewards. Thus, we show that computational modelling of affective dynamics can index the subjective value of intrinsic relative to extrinsic rewards, a ‘computational hedonometer’ that reflects both behavior and neural activity that quantifies the affective value of experience.

**SIGNIFICANCE STATEMENT:** Traditional economic indicators are increasingly recognized to provide an incomplete picture of what we value as a society. Standard economic approaches struggle to accurately assign values to non-market activities that nevertheless may be intrinsically rewarding, prompting a need for new tools to measure what really matters to individuals. Using a combination of neuroimaging and computational modeling, we show that despite their lack of instrumental value, intrinsic rewards influence subjective affective state and ventromedial prefrontal cortex activity. The relative degree to which extrinsic and intrinsic rewards influence affective state is predictive of their relative impacts on neural activity, confirming the utility of our approach for measuring the affective value of experiences and other non-market activities in individuals.

## INTRODUCTION

A key index of quality of life is subjective well-being which reflects “how people experience and evaluate their lives and specific domains and activities in their lives” (Oswald and Wu, 2010). Individuals with higher subjective well-being display lower mortality rates (Chida and Steptoe, 2008; Steptoe et al., 2015) and have a lower risk of disease (Davidson et al., 2010). In the workplace, employees who report higher subjective well-being have higher productivity without loss of output quality (Oswald et al., 2015), reduced rates of absenteeism (Pelled and Xin, 1999), and are rated more positively by their supervisors (Peterson et al., 2011). On this basis, maximizing subjective well-being should be of prime interest not only to individuals but also to companies and governments, as well as a target for health and economic policies (Dolan and White, 2007).

A problem arises when it comes to designing effective measures likely to increase well-being. When contemplating the future, people exhibit biases in *affective forecasting* when making predictions about what it would feel like to experience specific events, consistently misjudging how future events will impact their affective state and leading them to perform actions that may be detrimental to maximization of their subjective well-being (Wilson and Gilbert, 2005; Meyvis et al., 2010). In particular, people overestimate both the intensities and durations of their hedonic responses to future events, and this is referred to as an impact bias (Gilbert and Wilson, 2007; Morewedge and Buechel, 2013). Furthermore, the value of tangible goods can be quantified by prices or willingness-to-pay (Plassmann et al., 2007), but the value of intangible goods and experiences that are intrinsically rewarding (e.g., hobbies, recreational sports) are often more difficult to define or elicit accurately due to biases (Van de Mortel, 2008; Nisbet and Zelenski, 2011), while the predictive validity of implicit measures is unclear (Levesque et al., 2008; Keatley et al., 2013).

Neuroscience-informed methods can provide a means to evaluate the subjective value of an intrinsic reward (e.g., the experience of mastering a musical composition for its own sake), allowing extrinsic and intrinsic rewards to be compared using a common scale of objectively measured neural activity (FitzGerald et al., 2009). We hypothesized that extrinsic and intrinsic rewards would both influence affective states, and the extent of their relative influences should be reflected in regional brain activity. Recent studies (Rutledge et al., 2014, 2015; Vinckier et al., 2018) demonstrate that experience sampling during reward-based tasks can link affective and motivational responses to extrinsic reward. Here we extend this approach to investigate how affective state is influenced by the history of intrinsic rewards.

We developed a reinforcement learning task incorporating both an explicit reward component and a skilled performance component, where the latter did not affect payment (Figure 1A). On each trial, subjects selected one of two options, one of which was on average more rewarding than the other, and then navigated a cursor past a series of barriers (see Experimental Procedures). We hypothesized that the experience of successful skilled performance, a source of intrinsic reward, would influence the momentary happiness of subjects in a manner that is quantitatively akin to the impacts of extrinsic rewards and that this would also be evident at the level of neural activity.

**Figure 1.**
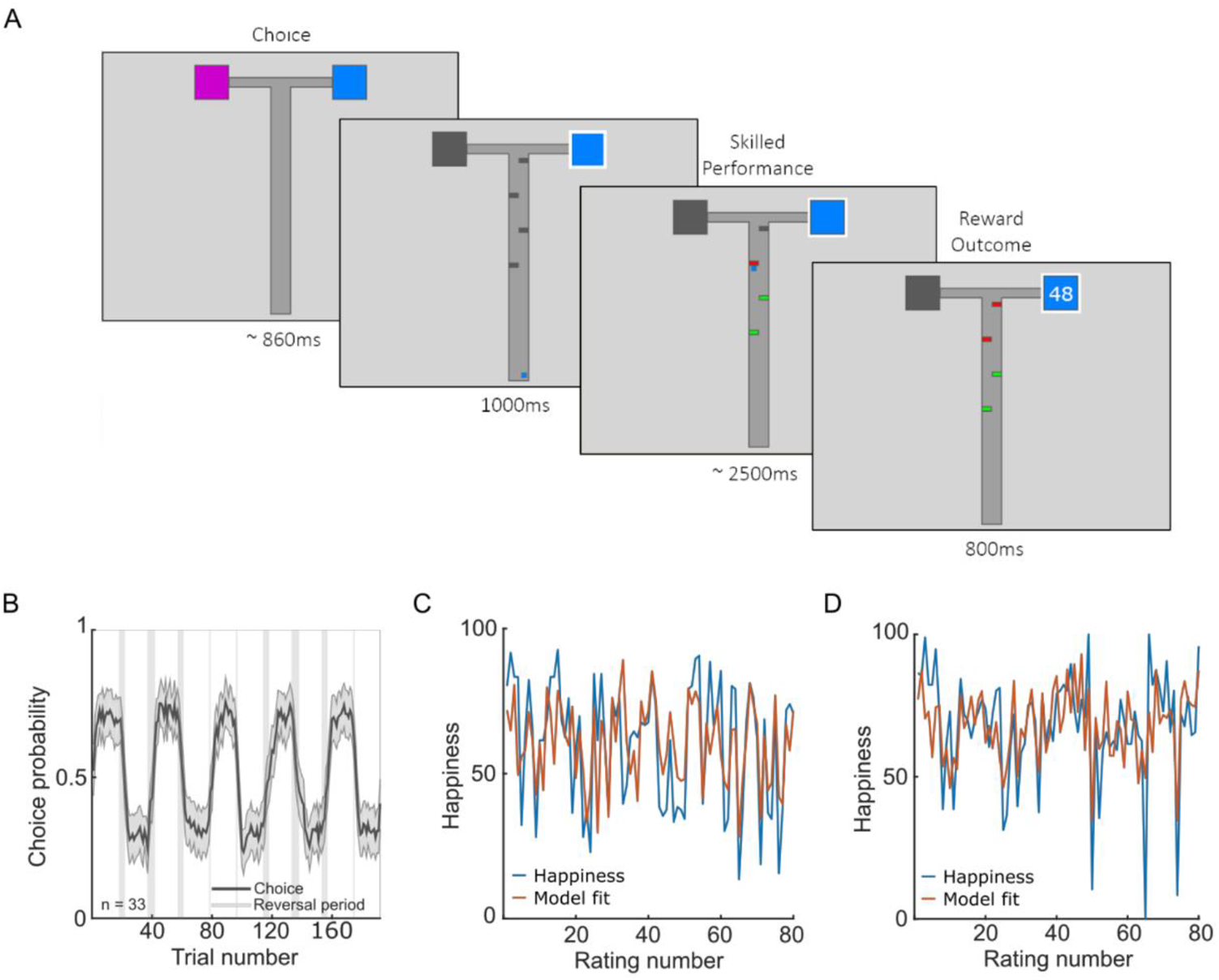
Extrinsic and intrinsic reward paradigm. (A) Subjects (n = 33) experienced both extrinsic and intrinsic rewards on each trial. A trial starts with subjects selecting from one or two available options each associated with an implicit extrinsic reward. One option on average leads to the larger reward (mean 50, SD 10) whereas the other leads to a lower reward (mean 25, SD 10) with a reversal every 19-23 trials. Four barriers then appear along the path to the outcome and a cursor appears at the bottom of the screen which automatically advances after a 1s delay. Subjects press left and right keys to navigate around barriers, constituting a form of skilled performance that can be intrinsically rewarding. Successfully avoiding a barrier turns it green whereas contact with a barrier turns it red. There is no financial penalty for contact with barriers nor financial benefit for avoiding them. Earnings depend only on the outcome delivered at the end of the trial. After every 2-3 trials, subjects report their current happiness by moving a cursor on a rating line. (B) Probability of choice to the initial high-reward option averaged across subjects (n = 33) in black. Shaded areas correspond to SEM. Grey vertical bands represent intervals where probability reversals could occur. (C, D) Happiness trajectories and model fits for a computational model with both reward and performance parameters are displayed for two example subjects (C: r^2^ = 0.45, D: r^2^ = 0.42). Also see Figure 2, Figure 3, Table 1 and Table 2.

## EXPERIMENTAL PROCEDURES

### Participants

37 healthy young adults (age: 25.8 ± 4.7, mean ± SD; 8 males, 29 females) were recruited through the University College London (UCL) Psychology Subject Database. Subjects were screened to ensure no history of neurological or psychiatric disorders. Four subjects were excluded due to excessive head movement during scanning, leaving a total of 33 subjects (age: 26.1 ± 4.9; 8 males, 25 females). The study was approved by the UCL research ethics committee, and all subjects gave written informed consent.

### Study Design

Subjects completed the experiment at the Wellcome Centre for Human Neuroimaging at UCL in an appointment that lasted approximately 90 minutes. Stimuli were presented in MATLAB (MathWorks, Inc.) using Cogent 2000. The layout of each trial resembled a T-Maze (Howe et al., 2013). On each trial, subjects selected a blue or magenta box, one of which resulted in 50 points on average and the other which resulted in 25 points on average. The standard deviation of points received for each box was 10. Points assigned based on draws from Gaussian distributions. Every 19-23 trials, a reversal occurred where the box that previously contained the higher number of points on average now contained a lower number of points and vice versa. On half of the trials, subjects were afforded a free choice. For the remaining half, subjects were only presented with a single option. After a choice was made, the chosen option was indicated and four barriers appeared on the screen along with a small cursor at the bottom of the screen. Following a 1s delay, the cursor automatically advanced along the path to the outcome. Subjects were able to control the horizontal position of the cursor to avoid colliding with barriers. If they passed a barrier without colliding with it, the barrier turned green. Contact with a barrier turned it red and provided immediate feedback about performance. Subjects then had to press the appropriate directional key to navigate around the barrier for the cursor to continue advancing on its course. Crucially, the subjects’ final payment depended only on the number of points accumulated across the experiment and not their ability to quickly navigate past barriers. After the cursor had entered the chosen box, the outcome was displayed for 800ms after a 1.5s delay. Total cumulative points were displayed on the top right of the screen throughout the experiment. Subjects were presented with the question, “How happy are you at this moment?” after every 2-3 trials. After a 1s delay period, a rating line appeared with a cursor at the midpoint and subjects had 4s to move a cursor along the scale with button presses. The left end of the line was labelled “very unhappy” and the right end of the line was labelled “very happy”.

### Staircase Procedure

To ensure that differences in affective responses were not due to skill-related differences in how often each subject collided with barriers, we used a standard staircase procedure called the Parametric Estimation by Sequential Testing (PEST) (Taylor and Creelman, 1967). This procedure calibrated the speed at which the cursor moved for every subject such that they did not contact the barriers on approximately 70% of trials. This calibration was carried out over 60 trials prior to the start of the task in the scanner. Continuation of the procedure during the task allowed small adjustments (e.g., to compensate for any fatigue) to maintain consistent successful skill performance.

### Questionnaire Measures

Subjects were administered the Beck Depression Inventory (BDI-II) (Beck et al., 1996), Apathy Evaluation Scale (AES) (Marin et al., 1991), and Apathy Motivation Index (AMI) (Ang et al., 2017).

### Image Acquisition

MRI scanning took place at the Wellcome Centre for Human Neuroimaging at UCL using a Siemens Prisma 3-Tesla scanner equipped with a 64-channel head coil. Functional images were acquired with a gradient echo T2*-weighted echo-planar sequence with whole-brain coverage. Each volume consisted of 48 slices with 3mm isotropic voxels [repetition time (TR): 3.36s; echo time (TE): 30ms; slice tilt: 0°] in ascending order. A field map [double-echo FLASH, TE1 = 10ms, TE2 = 12.46ms] with 3mm isotropic voxels (whole-brain coverage) was also acquired for each subject to correct the functional images for any inhomogeneity in magnetic field strength. Subsequently, the first 6 volumes of each run were discarded to allow for T1 saturation effects. Structural images were T1-weighted (1 × 1 × 1 mm resolution) images acquired using a MPRAGE sequence.

### Model-based Analyses

Models were fit to happiness ratings in individual subjects by minimizing the residual sum of squares between actual and predicted happiness ratings, and this also served as the objective function for the optimizer. Model fitting was performed using the *fmincon* optimizer in MATLAB (MathWorks, Inc). The significance for individual parameters was determined using likelihood ratio tests comparing the full model with a model that had only a reward or performance parameter but not both. The significance of those tests is indicated by filled circles in Figure 4. Note that models were first fit to the raw happiness ratings in order to test the relationship between the happiness baseline mood parameter (denoted w0 in the equations below) and questionnaire measures to replicate findings in the literature. Models were then fit to standardized ratings. Normalizing ratings prevents individuals with greater variance in their ratings from having a disproportionate effect on model comparisons. The standard deviation of ratings differs widely across participants although rating variance is known to be stable in time (Rutledge et al., 2015) and across tasks (Blain and Rutledge, 2020).

### Recovery Analysis

To ensure that the model parameters were recoverable, we performed model recovery and parameter recovery analyses following established procedures (Wilson & Collins). To test for parameter recovery, we first estimated the parameters for each participant. Then, we simulated data with each of the four generative models using parameters estimated for each participant. We added Gaussian noise in the simulations based on individual-level noise estimates. We then estimated parameters from the generated data using the same procedure as applied to the actual mood dynamics data (n = 33).

## RESULTS

Subjects completed two trial blocks while in the MRI scanner. We first asked whether subjects could learn the reward contingencies (Figure 1B) and found that they could, making 85.8 ± 1.0%% (mean ± SEM, z = 5.0, p < 10^−6^) of choices to the current high-reward option. Subjects were not penalized for contact with barriers, and thus actual performance was non-instrumental to the receipt of eventual monetary reward. We observed no correlation between earnings and how often subjects successfully avoided barriers (ρ(31) = 0.21, p = 0.24). During debriefing, all 33 subjects reported that they believed there was no association between successful skilled performance and earnings.

Reports of affective state for example subjects are included in Figure 1C. On average, subjects reported being happier after receiving outcomes from the high-compared to low-reward option (high-reward:63.8 ± 1.9, low-reward: 59.5 ± 2.1, z = 4.7, p < 10^−5^), consistent with previous research (Rutledge et al., 2014, 2015). On average, subjects reported also being happier when they navigated through the barriers without collisions compared to when they contacted at least one barrier (without collisions: 63.5 ± 1.9; collision: 60.0 ± 2.1, z = 4.6, p < 10^−5^), suggesting that intrinsic rewards related to performance influence subjective affective state.

Because participants vary in how they use the scale, we next z-scored happiness ratings. Consistent with analyses using non-normalized ratings, subjects reported greater average happiness after receiving high compared to low rewards (high-reward: 0.08 ± 0.01, low-reward: −0.18 ± 0.02, z = 4.8, p < 10^−5^, Figure 2A). Subjects also reported being happier after navigating through the maze without contacting any barriers compared to when they collided with at least one barrier (without collisions: 0.08 ± 0.01; collision: −0.17 ± 0.03, z = 4.7, p < 10^−5^, Figure 2A), consistent with an impact of intrinsic rewards. There was considerable variation across subjects in terms of how much extrinsic rewards and skilled performance contributed to momentary happiness (Figure 2B), but there was no relationship between happiness for reward outcomes and happiness for skilled performance (ρ(31) = −0.20, p = 0.26).

**Figure 2.**
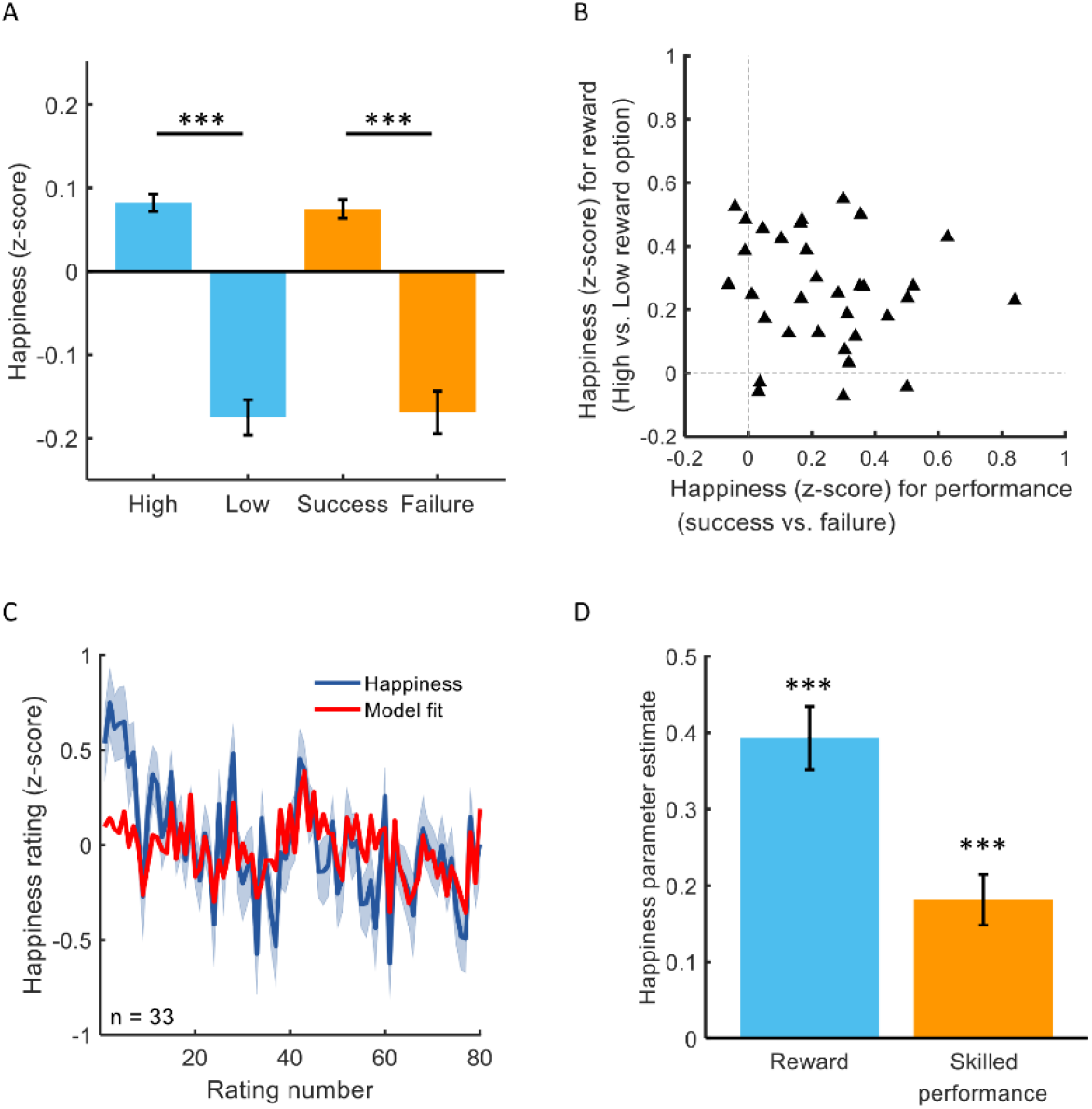
Computational modelling of affective dynamics. (A) Subjects were happier when they received a reward from high-compared to low-reward options (Z = 4.7, p < 10^−5^, in blue). Subjects were happier on average when they navigated through the barriers without contacting them, compared to when they contacted at least one barrier (Z = 4.6, p < 10^−5^, in orange). *** p < 0.001. (B) The majority of subjects (29 of 33) were happier after receiving a reward from a high-compared to low-reward option. The majority of subjects (29 of 33) were happier after successful compared to unsuccessful performance. There was no relationship between happiness for reward outcomes and happiness for skilled performance (ρ(31) = −0.20, p = 0.26). (C) Average happiness across all subjects and model fit is displayed for the computational model (n = 33, mean r^2^ = 0.26). (D) According to the computational model, happiness was significantly related to the history of extrinsic rewards in the form of points converted to money (Z = 4.9, p < 10^−5^) and also to the history of skilled performance, a proxy for intrinsic rewards (Z = 4.4, p < 10^−4^). *** p < 0.001.

### Computational model of affective dynamics

We next employed a previously established methodology (Rutledge et al., 2014, 2015; Blain and Rutledge, 2020) to quantify the extent to which rewards impacted on the affective state of our participants. In particular, we aim to replicate that (1) the recent history of reward influences happiness and (2) that the baseline happiness parameter correlates with depressive symptoms. To that end, we fit the raw happiness ratings. We considered influences that decay exponentially in time:

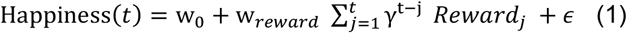

where *t* and *j* are trial numbers, w_0_ is a baseline mood parameter, w_reward_ captures the influence of reward which is the z-scored reward outcome of the selected option on each trial, and 0 ≤ ɣ ≤ 1 represents a forgetting factor that reduces the impact of distal relative to recent events. If this parameter is equal to 0, only the most recent reward outcome influences happiness. The model includes a Gaussian noise term, ϵ ∼ N(0, σ) The parameters of this model are recoverable (see Figure 3A and Table 1 for details about parameter recovery). Parameters were first fit to non-normalized happiness ratings in each individual subject. The mean r^2^ was 0.26 ± 0.03 and the mean forgetting factor was 0.40 ± 0.06 (mean ± SEM, Figure 1C for example subjects). Consistent with previous findings (Rutledge et al., 2014, 2015), happiness was significantly associated with the history of reward (w_reward_ = 0.06 ± 0.01; Wilcoxon signed rank test: z = 4.7, p < 10^−5^). Sigma was estimated to be on average 0.13 ± 0.01.

**Figure 3.**
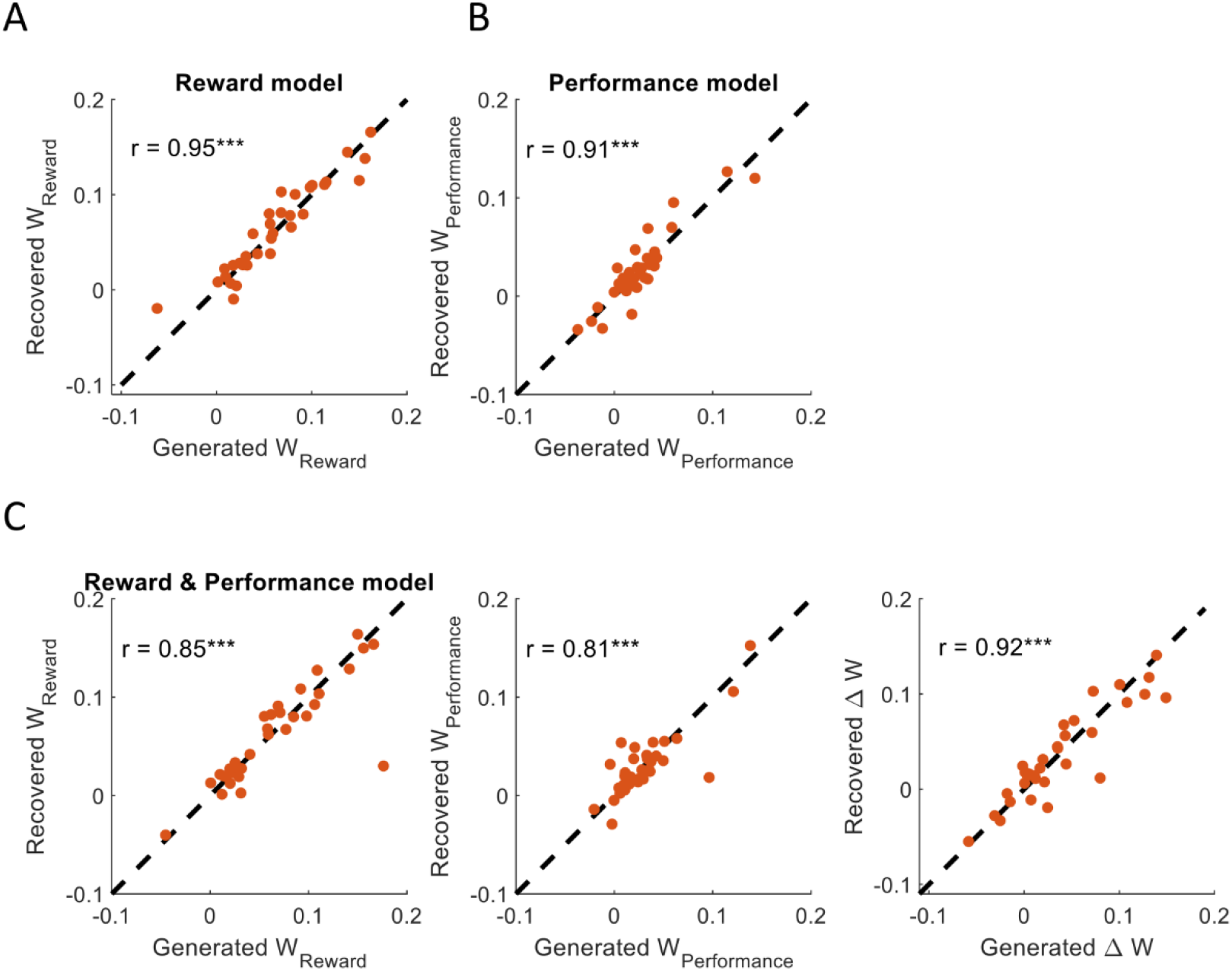
Parameter recovery analysis for the reward model (A), the performance model (B), and the reward and performance model (C). The X-axis represents the parameter values used to generate the data and the Y-axis corresponds to the estimated parameters. The parameters are recoverable with no bias. See Experimental Procedures for details. *** P < 10-7

**Table 1.** Model parameter recovery results.

| Model | Spearman $\rho$ between generated and estimated parameters | | | | |
| --- | --- | --- | --- | --- | --- |
| | $W_0$ | $W_{\text{reward}}$ | $W_{\text{performance}}$ | $\gamma_1$ | $\gamma_2$ |
| Reward | 0.99*** | 0.94 *** | - | 0.55** | - |
| Performance | 0.99*** | - | 0.88*** | 0.4* | - |
| Reward and performance | 0.98*** | 0.86 *** | 0.86*** | 0.84*** | - |
| Reward and performance (separate $\gamma$) | 0.99*** | 0.68*** | 0.88*** | 0.50** | 0.73*** |

The values correspond to the Spearman correlation between the generated parameters and the estimated parameters of 33 agents. See Experimental Procedures for details. \* $p < 0.05$ , \*\* $p < 0.01$ , \*\*\* $p < 0.001$

Likewise, consistent with previous findings during risky decision making (Rutledge et al., 2017), we found that baseline mood parameters, estimated using raw happiness ratings while accounting for mood dynamics due to reward history, were negatively correlated with symptom severity assessed using the Beck Depression Inventory (BDI-II; Beck et al., 1996; Spearman ρ(31) = −0.35, p = 0.046). This result shows that depressive symptoms relate to happiness ratings during a novel task including a performance component consistent with previous findings during risky decision making (Rutledge et al., 2017) and learning in volatile environments (Blain and Rutledge, 2020). This relationship is consistent with an affective set point, which happiness returns to over time, that is lower in individuals with a greater symptom load.

We also found baseline mood parameters tended to be negatively related apathy as measured by Apathy Evaluation Scale (AES) (Marin et al., 1991) (ρ(31) = −0.32, p = 0.07) and behavioral apathy as assessed by the Apathy Motivation Index (AMI) (27) (ρ(31) = −0.33, p = 0.06; see Table 2). The first happiness rating before the start of the first trial was positively correlated with baseline mood parameter (ρ(31) = 0.46, p = 0.007). In contrast to baseline mood parameters, first happiness ratings were not significantly correlated with BDI-II (ρ(31) = −0.21, p = 0.25) or AES (ρ(31) = −0.17, p = 0.35), but was correlated with behavioral AMI (ρ(31) = −0.39, p = 0.027). We found no correlation between baseline mood parameter and the average staircased cursor speed (ρ(31) = −0.01, p = 0.95), suggesting that the speed of the cursor was not associated with persistent affective state.

**Table 2.** Model comparison results.

| Model | Parameters | Mean $r^2$ | BIC | ΔBIC |
| --- | --- | --- | --- | --- |
| Reward | 2 | 0.19 | -326 | 145 |
| Performance | 2 | 0.09 | -26 | 445 |
| Reward and Performance | 3 | 0.26 | -471 | 0 |
| Reward and Performance (separate $\gamma$) | 4 | 0.27 | -351 | 120 |

Bayesian Information Criterion (BIC) scores are summed across 33 subjects. The winning model (lowest BIC) was the model with both reward and performance having the same forgetting factor $\gamma$ rather than a model where the influence of past reward and performance differs in their forgetting factor. $\Delta$ BIC refers to the difference in BIC between each model and the winning model. Ratings are z-scored to prevent individuals with greater rating variance from disproportionately influencing model comparison.

We next z-scored happiness ratings to better evaluate the relative contributions of extrinsic and intrinsic reward to affective state. To that end, we z-scored the happiness ratings, thereby preventing individuals with greater rating variance from disproportionally affecting analyses. With happiness ratings centered on zero, as well as Rewards and Performance vectors, any constant term would be expected to be near zero and we omitted the w0 from analyses with z-scored ratings. We expanded the model to include an additional term that accounts also for influences pertaining to skilled performance:

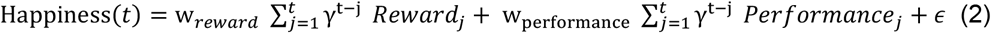

where *t* and *j* are trial numbers, w_reward_ and w_performance_ capture the influence of task events related to reward and performance, respectively, and 0 ≤ ɣ ≤ 1 represents a forgetting factor that reduces the impact of distal relative to recent events. The model includes a Gaussian noise term, ϵ ~ N(0, σ). The model parameters were indeed recoverable (see Figure 2C and table 2 and methods for details). Reward is the z-scored outcome of the selected option on each trial, and performance is the z-scored result of whether a barrier was contacted on each trial, assigning a 1 when no barriers were contacted and 0 if at least one barrier was contacted. This simple model explained a substantial amount of variance in happiness with r^2^ = 0.26 ± 0.03 (mean ± SEM, Figure 2C). Weights for both performance (w_performance_ = 0.18 ± 0.03; z = 4.4, p < 10^−4^, Figure 2D) and reward (w_reward_ = 0.39 ± 0.04, z = 4.9, p < 10^−5^, Figure 2D) were positive on average. The forgetting factor ɣ was 0.48 ± 0.05 (mean ± SEM), indicating that happiness depended on the past 4^-5^ trials on average. Sigma was estimated to be on average 0.85 ± 0.02.

In previous studies we found expectations of reward exerted a substantial influence on happiness (Rutledge et al., 2014, 2015; Blain and Rutledge, 2020). In the current study, we used high- and low-reward distributions with minimal overlap to maximize learning accuracy. We also employed a staircase to keep skilled performance stable and at a similar level across individuals. These features render the current design unsuitable for quantifying the impact of expectations on happiness. We chose a design that maximized our power for quantifying individual differences in the relative subjective values of extrinsic and intrinsic rewards.

Model comparison (Table 2) shows that a model with parameters for past rewards and performance (mean r^2^ = 0.26) outperformed models containing individual terms for reward (mean r^2^ = 0.19) or performance (mean r^2^ = 0.09) alone. These results show that the happiness of subjects in this task is, on average, dependent on both receipt of explicit rewards (e.g., money) and the non-instrumental experience of skilled performance.

We found considerable variation across individuals in how much reward outcomes contributed to affective dynamics, even though subjects on average learned reward contingencies to a similar degree (Figure 4A). Despite performance being held constant due to staircasing of cursor speed (successful performance: 69.1 ± 2.4%, mean ± SD, Figure 4B), there was considerable variation also across individuals in how much non-instrumental performance influenced affective state. Many subjects showed a negligible impact of successful performance on affective state, despite a similar level of successful performance. Furthermore, learning choice accuracy was not correlated with either happiness reward parameters (ρ(31) = 0.12, p = 0.49) or successful skilled performance (ρ(31) = −0.05, p = 0.78).

**Figure 4.**
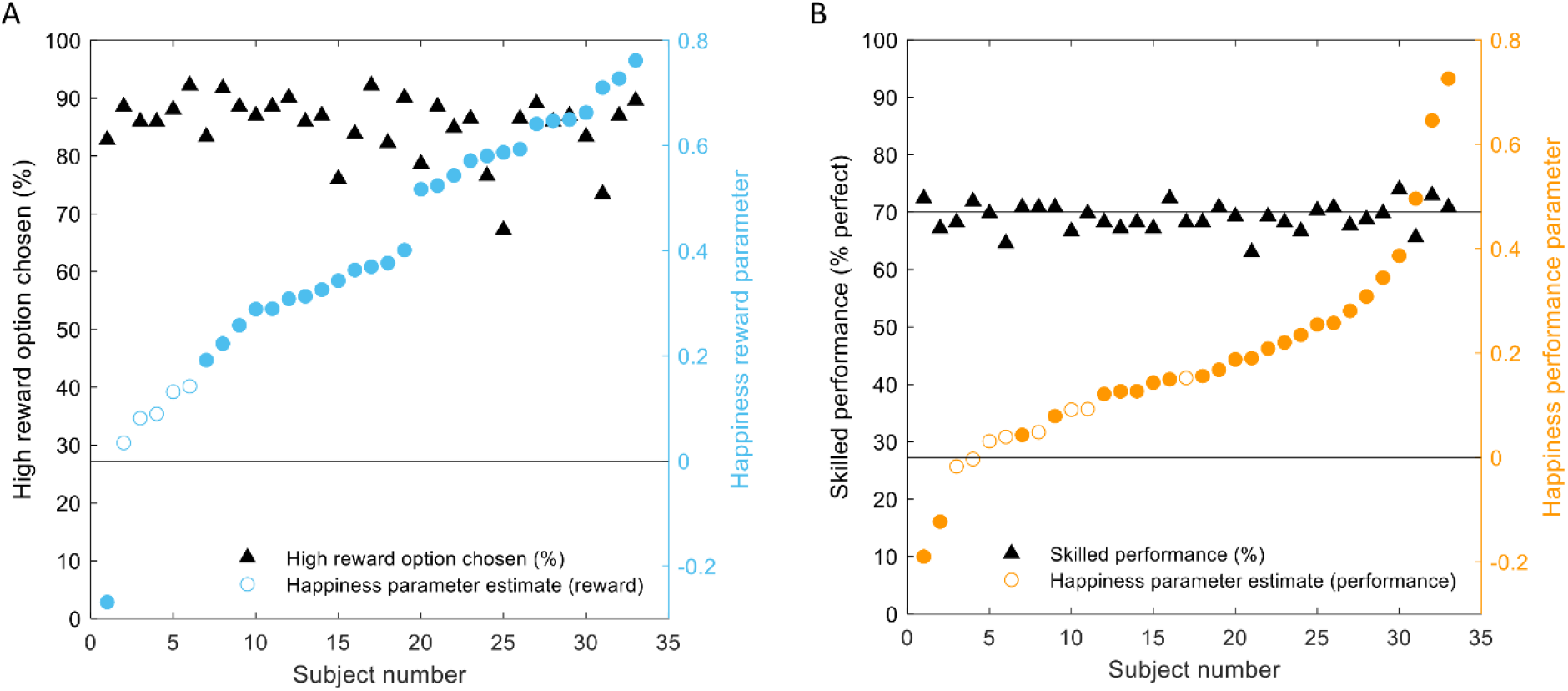
Computational model parameters and task behavior. (A, B) The contribution of reward to happiness varied across subjects despite a similar high choice accuracy across subjects. Despite titrating difficulty at the individual level to match performance across subjects at 70%, subjects displayed considerable variation in the degree to which performance impacted affective state as captured by the computational model. Filled circles indicate betas that are significant at the individual level.

Intrinsic rewards can be associated with an increased motivation or metacognitive strategy to improve performance over time (Son and Metcalfe, 2005). Prior to scanning, participants completed 60 practice trials to determine an appropriate starting speed for the experiment. W_performance_ was positively correlated with the starting cursor speed (ρ(31) = 0.38, p = 0.03). There was no correlation between percent successful skilled performance and w_performance_ derived from the happiness model (ρ(31) = 0.056, p = 0.76). Intrinsic rewards are often thought as resulting from uncertainty reduction, or from learning progress (Gottlieb and Oudeyer, 2018). However, we did not find any significant difference in the median cursor speed between blocks (z = 0.63, p = 0.53), suggesting that participants were at a stable level of performance from the start that did not improve over time. Similarly, w_performance_ was not significantly different between blocks (z = 1.47, p = 0.14). These results together suggest that performing this task accurately was intrinsically rewarding with a stable relationship between performance and happiness despite no signs of learning progress during the experiment.

We then checked whether we can extend the link between the baseline mood parameter from the reward model (see above) and apathy and depression scores to the baseline mood parameter of models including a performance term. Results indicate a trend towards the same relationship as for the reward model (see Table 3).

**Table 3.** Correlation between baseline mood parameter and questionnaire score. Values correspond to the Spearman coefficient ρ. *p < 0.05, ⴕ < 0.1

| | $W_0$ reward | $W_0$ performance | $W_0$ reward & performance |
| --- | --- | --- | --- |
| *BDI* | -0.35* | -0.31 † | -0.34† |
| *AES* | -0.32 † | -0.32 † | -0.30 † |
| *bAMI* | -0.33 † | -0.32 † | -0.29 † |

### Neural correlates of extrinsic and intrinsic rewards

Having established inter-individual variability in the impact of outcomes and performance on reported happiness, we next asked whether this variability was also predictive of neural responses to both rewards and performance. The experiment was separated into two scans and we first evaluated whether happiness model parameters were stable across scans. We found that both extrinsic (ρ(33) = 0.35, p = 0.044) and intrinsic (ρ(33) = 0.35, p = 0.044) reward computational parameters were positively correlated across the two scans.

We regressed event-related activity on parametrically modulated task events to assess brain activity related to receipt of extrinsic and intrinsic rewards. We found an effect of reward magnitude at time of outcome in vmPFC (Figure 4A, top: −3, 38, −1; t_32_ = 5.92, p < 0.05 Family-Wise-Error (FWE) cluster-corrected at the whole brain level), as well as an effect of successful skilled performance in an overlapping region of the vmPFC (Figure 4A, bottom: −3, 50, −1; t_32_ = 4.24, p < 0.05 FWE cluster-corrected).

The vmPFC is widely implicated in representation of subjective reward value. On this basis, we used an independent vmPFC mask from a meta-analysis of subjective value studies of extrinsic reward for further analysis (Bartra et al., 2013). Within this region-of-interest (ROI), we extracted weights for reward magnitude and skilled performance from each individual subject. We found that within this independent ROI, BOLD activity was significantly associated with both reward magnitude (0.26 ± 0.08, Z = 3.0, p = 0.0029) and skilled performance (0.38 ± 0.13, Z = 2.8, p = 0.0052, Figure 5B).

**Figure 5.**
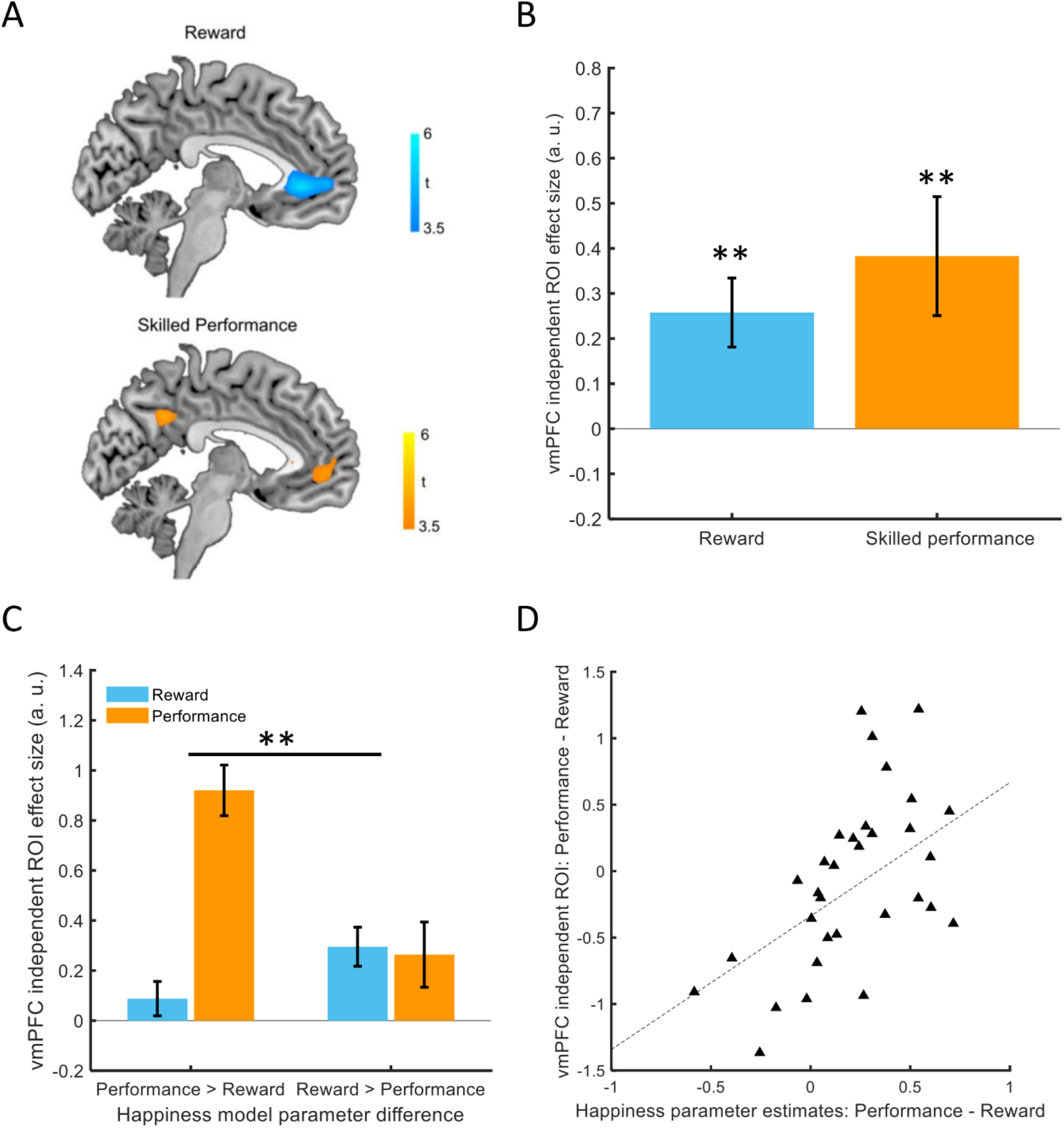
Relative affective impacts of reward and performance predict vmPFC activity. (A) *Top*. BOLD activity in vmPFC was parametrically modulated by reward magnitude (Peak: −3, 38, −1). *Bottom*. Bold activity in an overlapping region of vmPFC was modulated by trial-by-trial successful skilled performance (Peak: −3, 50, −1). (B) An independent vmPFC ROI shows modulation by both reward magnitude and skilled performance (both p < 0.01). (C) In the independent vmPFC ROI, subjects with higher performance than reward weights in the computational analysis of affective dynamics displayed stronger neural responses in the vmPFC for performance than subjects with higher reward than performance weights (p = 0.003). (D) The difference between performance and reward weights in the happiness computational model predicted the difference in vmPFC neural responses for successful skilled performance relative to reward magnitude (ρ(31) = 0.50, p = 0.003). * p < 0.05, ** p < 0.01.

Having established that neural responses in vmPFC are associated with both extrinsic and intrinsic rewards, we next examined whether neural responses were predicted by computational parameters estimated from individual affective dynamics. Across subjects, we found a positive relationship (ρ(31) = 0.50, p = 0.003, Figure 5D) between the relative weights for extrinsic and intrinsic rewards in our happiness computational model and the relative effect sizes for neural responses in the vmPFC. Initial happiness ratings deviate from model predictions on average (Figure 2C). The relationship between relative happiness weights and relative neural effect sizes was still present after removing the initial 10% of ratings (ρ(31) = 0.54, p = 0.0015). The relationship was also present after removing the initial 10% and detrending the remaining ratings before estimating model parameters (ρ(31) = 0.49, p = 0.0038).

We also subdivided subjects into two groups comprising a group with higher W_performance_ than reward parameters and a group with the opposite pattern. The group with higher performance than reward parameters showed greater vmPFC responses for skilled performance compared to the group with larger reward than performance parameters (Z = 2.8, p = 0.0047, Figure 5C). These findings suggest that the pattern of momentary affective dynamics reflects the impact of both extrinsic and intrinsic rewards and is mirrored at the level of vmPFC activity.

## DISCUSSION

Using experience sampling (Reis and Gable, 2000; Kahneman et al., 2004) combined with functional neuroimaging, we show that extrinsic and intrinsic rewards contribute to affective dynamics (i.e., happiness). Recent studies demonstrate that computational approaches can quantify consistent relationships between subjective feelings and value-based decision making (Rutledge et al., 2014; Eldar et al., 2016, 2018; Vinckier et al., 2018; Blain and Rutledge, 2020), including in relation to individual social preferences (Rutledge et al., 2016). Here, using the same computational approach applied during reinforcement learning, we show that momentary happiness is influenced by both extrinsic and intrinsic rewards. The computational parameters we extract from affective dynamics enabled us to quantify, within a common value scale, the relative affective value of intrinsic relative to extrinsic rewards. Our key finding here is that the relative weight of intrinsic and extrinsic reward extracted from affective dynamics predicts neural activity in the vmPFC, a region proposed to represent rewards in a common neural currency (Chib et al., 2009; Levy and Glimcher, 2011, 2012), validating our computational approach.

While improvements in skilled performance can be enhanced by rewarding individuals for performance (Sugawara et al., 2012), holding performance constant across subjects allowed us to investigate how happiness varied independently of the level of skill individuals manifest in the task. We show that individuals, whose happiness was substantially influenced by intrinsic rewards, had increased vmPFC BOLD responses for successful versus unsuccessful skilled performance, relative to individuals whose happiness was influenced more by extrinsic rewards.

The vmPFC is known to represent the value of different types of goods, including food and juice (Padoa-Schioppa, 2007; Hare et al., 2011), money (De Martino et al., 2006), aesthetic judgments (Kawabata and Zeki, 2004; Jacobsen et al., 2006), and even perceived pleasantness (Plassmann et al., 2008). This suggests that vmPFC plays a central role in representing qualitatively different types of goods on a common scale, an operation that can facilitate making decisions between otherwise incommensurable goods (Chib et al., 2009; Levy and Glimcher, 2011, 2012). Our study builds on these prior results by now identifying an association between vmPFC BOLD activity and intrinsic rewards, here the experience of performing a skilled task without error. Whole-brain analysis showed that the representation of subjective intrinsic reward values involved an adjacent region in the vmPFC, anterior to the representation for extrinsic rewards but still residing within a central vmPFC cluster (Clithero and Rangel, 2014), a finding that parallels a distinction between experienced and decision values previously mapped to anterior and posterior vmPFC, respectively (Smith et al., 2010).

The vmPFC has been demonstrated to play a role in affect with subjective emotional experiences elicited by images and pleasurable music leading to changes in both vmPFC BOLD activity and regional cerebral blood flow (Blood and Zatorre, 2001; Zald et al., 2002; Winecoff et al., 2013). Damage to the vmPFC can lead to aberrant emotional responses (Koenigs et al., 2007; Zald and Andreotti, 2010; Hiser and Koenigs, 2018) and maladaptive decision making in environments where emotional regulation may be useful (Grossman et al., 2010; Spaniol et al., 2019). Numerous studies suggest that subjective reward values are represented by vmPFC neural activity. Unfortunately, the constraints and expense of neuroimaging makes it impractical as an every-day tool for assessing individual values for non-market activities. The strong association between neural responses for intrinsic and extrinsic rewards and computational parameters extracted from affective dynamics suggests that computational models combined with experience sampling can provide a valid measure for the subjective reward value of experience.

A limitation of the current study is that the staircase procedure we used does not allow us to address questions related to the intrinsic motivation for learning of our subjects. The staircase procedure can be useful for study of interindividual variation either by keeping performance constant across individuals despite differences in abilities (Fleming et al., 2010) or for tailoring choice options to individuals (Klein-Flügge et al., 2015). Using the staircase procedure meant that subjects quickly reached the limit by which they could improve performance. Our design is thus unsuitable for studying intrinsic motivation pertaining to learning. However, such a framework for measuring affective value could be valuable for other features related to intrinsic rewards (Blain and Sharot, 2021), like metacognitive control and learning (Son and Sethi, 2006), resource allocation under external pressures (Son and Metcalfe, 2005), as well as curiosity-driven exploration of the environment where rewards may be more dependent on the learning progress of an individual (Gottlieb and Oudeyer, 2018).

Humans exhibit biases when it comes to predicting how future events are likely to impact on their affective states, and are prone to making sub-optimal decisions by misjudging the hedonic consequences of options (Wilson and Gilbert, 2005; Meyvis et al., 2010; Nisbet and Zelenski, 2011). Increasing subjective well-being is widely believed to be an appropriate societal goal (OECD, 2020), but these biases pose a difficulty for enacting policies that are likely to be successful. Additional factors such as social desirability bias (Van de Mortel, 2008) can decrease the reliability of self-reported values when an individual’s assessment of a hypothetical experience or good, such as the availability of public parks, differs from prevailing social norms. An advantage of our method is that it can be in principle applied to any repeatable experience without a need to probe people explicitly about the content of those experiences, reducing biases associated with social desirability. For example, affective dynamics reflect depressive symptoms (Rutledge et al., 2017; Blain and Rutledge, 2020), show consistent relationships to reward in the lab and outside the lab in anonymous participants who did not interact with an experimenter (Rutledge et al., 2014), and allow quantification of the extent of guilt and envy in response to social inequality (Rutledge et al., 2016). A potential application of our approach, yet to be tested, would be to combine our computational approach with experience sampling in different naturalistic settings such as a corporate workplace, in order to identify factors important for employee well-being. Thus, the approach we use in this study demonstrates a novel tool for understanding preferences and well-being.

Over a century ago, Francis Edgeworth described an idealized instrument, which he called a hedonometer, for ‘continually registering the height of pleasure experienced by an individual’ (Edgeworth, 1881). Here, we introduce a ‘computational hedonometer’ that has a distinct advantage over Edgeworth’s hypothetical hedonometer in that it mathematically quantifies the relative contributions of different factors to an affective state, including the relative values of intrinsic and extrinsic rewards. We validate our computational tool using objective neural measurements, suggesting that computational parameters can capture the affective values for abstract goods and experiences that may be otherwise challenging to accurately quantify.

## ACKNOWLEDGMENTS

We thank Tobias Hauser, Matilde Vaghi, and Rachel Bedder for helpful comments. B.C. is a predoctoral fellow of the International Max Planck Research School on Computational Methods in Psychiatry and Ageing Research. The participating institutions are the Max Planck Institute for Human Development and the University College London (UCL). B.C. is also supported by a scholarship from the Singapore Institute of Management. R.J.D. holds a Wellcome Trust Investigator Award (098362/Z/12/Z). R.B.R. is supported by a Medical Research Council Career Development Award (MR/N02401X/1), a 2018 NARSAD Young Investigator Grant (27674) from the Brain and Behavior Research Foundation, P&S Fund, and by the National Institute of Mental Health (1R01MH124110). The Max Planck UCL Centre is a joint initiative supported by UCL and the Max Planck Society. The Wellcome Centre for Human Neuroimaging is supported by core funding from the Wellcome Trust (203147/Z/16/Z).

## REFERENCES

Ang Y-S, Lockwood P, Apps MA, Muhammed K, Husain M (2017) Distinct subtypes of apathy revealed by the apathy motivation index. PloS one 12:e0169938.

Bartra O, McGuire JT, Kable JW (2013) The valuation system: A coordinate-based meta-analysis of BOLD fMRI experiments examining neural correlates of subjective value. NeuroImage 76:412–427.

Beck AT, Steer RA, Ball R, Ranieri WF (1996) Comparison of beck depression inventories-ia and-ii in psychiatric outpatients. Journal of Personality Assessment 67:588–597.

Blain B, Rutledge RB (2020) Momentary subjective well-being depends on learning and not reward Lee D, ed. eLife 9:e57977.

Blain B, Sharot T (2021) Intrinsic reward: potential cognitive and neural mechanisms. Current Opinion in Behavioral Sciences 39:113–118.

Blood AJ, Zatorre RJ (2001) Intensely pleasurable responses to music correlate with activity in brain regions implicated in reward and emotion. PNAS 98:11818–11823.

Chib VS, Rangel A, Shimojo S, O’Doherty JP (2009) Evidence for a common representation of decision values for dissimilar goods in human ventromedial prefrontal cortex. Journal of Neuroscience 29:12315–12320.

Chida Y, Steptoe A (2008) Positive psychological well-being and mortality: a quantitative review of prospective observational studies. Psychosom Med 70:741–756.

Clithero JA, Rangel A (2014) Informatic parcellation of the network involved in the computation of subjective value. Soc Cogn Affect Neurosci 9:1289–1302.

Davidson KW, Mostofsky E, Whang W (2010) Don’t worry, be happy: positive affect and reduced 10-year incident coronary heart disease: the Canadian Nova Scotia Health Survey. Eur Heart J 31:1065–1070.

De Martino B, Kumaran D, Seymour B, Dolan RJ (2006) Frames, Biases, and Rational Decision-Making in the Human Brain. Science 313:684–687.

Dolan P, White MP (2007) How Can Measures of Subjective Well-Being Be Used to Inform Public Policy? Perspect Psychol Sci 2:71–85.

Edgeworth FY (1881) Mathematical psychics: An essay on the application of mathematics to the moral sciences. Kegan Paul.

FitzGerald THB, Seymour B, Dolan RJ (2009) The Role of Human Orbitofrontal Cortex in Value Comparison for Incommensurable Objects. J Neurosci 29:8388–8395.

Fleming SM, Weil RS, Nagy Z, Dolan RJ, Rees G (2010) Relating Introspective Accuracy to Individual Differences in Brain Structure. Science 329:1541–1543.

Gilbert DT, Wilson TD (2007) Prospection: experiencing the future. Science 317:1351–1354.

Gottlieb J, Oudeyer P-Y (2018) Towards a neuroscience of active sampling and curiosity. Nature Reviews Neuroscience 19:758–770.

Grossman M, Eslinger PJ, Troiani V, Anderson C, Avants B, Gee JC, McMillan C, Massimo L, Khan A, Antani S (2010) The role of ventral medial prefrontal cortex in social decisions: converging evidence from fMRI and frontotemporal lobar degeneration. Neuropsychologia 48:3505–3512.

Hare TA, Malmaud J, Rangel A (2011) Focusing Attention on the Health Aspects of Foods Changes Value Signals in vmPFC and Improves Dietary Choice. J Neurosci 31:11077–11087.

Hiser J, Koenigs M (2018) The multifaceted role of ventromedial prefrontal cortex in emotion, decision-making, social cognition, and psychopathology. Biol Psychiatry 83:638–647.

Howe MW, Tierney PL, Sandberg SG, Phillips PE, Graybiel AM (2013) Prolonged dopamine signalling in striatum signals proximity and value of distant rewards. nature 500:575–579.

Jacobsen T, Schubotz R, Höfel L, Cramon Y (2006) Brain correlates of aesthetic judgment of beauty. Neuroimage 29:276–285.

Kahneman D, Krueger AB, Schkade DA, Schwarz N, Stone AA (2004) A survey method for characterizing daily life experience: The day reconstruction method. Science 306:1776–1780.

Kawabata H, Zeki S (2004) Neural Correlates of Beauty. Journal of Neurophysiology 91:1699–1705.

Keatley D, Clarke DD, Hagger MS (2013) The predictive validity of implicit measures of self-determined motivation across health-related behaviours. British Journal of Health Psychology 18:2–17.

Klein-Flügge MC, Kennerley SW, Saraiva AC, Penny WD, Bestmann S (2015) Behavioral Modeling of Human Choices Reveals Dissociable Effects of Physical Effort and Temporal Delay on Reward Devaluation. PLOS Computational Biology 11:e1004116.

Koenigs M, Young L, Adolphs R, Tranel D, Cushman F, Hauser M, Damasio A (2007) Damage to the prefrontal cortex increases utilitarian moral judgements. Nature 446:908–911.

Levesque C, Copeland KJ, Sutcliffe RA (2008) Conscious and nonconscious processes: Implications for self-determination theory. Canadian Psychology/Psychologie canadienne 49:218.

Levy DJ, Glimcher PW (2011) Comparing apples and oranges: using reward-specific and reward-general subjective value representation in the brain. J Neurosci 31:14693–14707.

Levy DJ, Glimcher PW (2012) The root of all value: a neural common currency for choice. Current Opinion in Neurobiology 22:1027–1038.

Marin RS, Biedrzycki RC, Firinciogullari S (1991) Reliability and validity of the Apathy Evaluation Scale. Psychiatry research 38:143–162.

Meyvis T, Ratner RK, Levav J (2010) Why don’t we learn to accurately forecast feelings? How misremembering our predictions blinds us to past forecasting errors. J Exp Psychol Gen 139:579–589.

Morewedge CK, Buechel EC (2013) Motivated underpinnings of the impact bias in affective forecasts. Emotion 13:1023–1029.

Nisbet EK, Zelenski JM (2011) Underestimating nearby nature: Affective forecasting errors obscure the happy path to sustainability. Psychological science 22:1101–1106.

OECD O (2020) How’s Life? 2020 : Measuring Well-being | OECD iLibrary. Available at: ../sdd-2020-25-en/index.html [Accessed March 22, 2021].

Oswald AJ, Proto E, Sgroi D (2015) Happiness and Productivity. Journal of Labor Economics 33:789–822.

Oswald AJ, Wu S (2010) Objective Confirmation of Subjective Measures of Human Well-Being: Evidence from the U.S.A. Science 327:576–579.

Padoa-Schioppa C (2007) Orbitofrontal cortex and the computation of economic value. Ann N Y Acad Sci 1121:232–253.

Pelled LH, Xin KR (1999) Down and Out: An Investigation of the Relationship between Mood and Employee Withdrawal Behavior. Journal of Management 25:875–895.

Peterson S, Luthans F, Avolio BJ, Walumbwa FO, Zhang Z (2011) Psychological capital and employee performance: A latent growth modeling approach. Personnel Psychology 64:427–450.

Plassmann H, O’Doherty J, Rangel A (2007) Orbitofrontal cortex encodes willingness to pay in everyday economic transactions. J Neurosci 27:9984–9988.

Plassmann H, O’Doherty J, Shiv B, Rangel A (2008) Marketing actions can modulate neural representations of experienced pleasantness. PNAS 105:1050–1054.

Reis HT, Gable SL (2000) Event-sampling and other methods for studying everyday experience.

Rutledge RB, de Berker AO, Espenhahn S, Dayan P, Dolan RJ (2016) The social contingency of momentary subjective well-being. Nat Commun 7 Available at: http://www.ncbi.nlm.nih.gov/pmc/articles/PMC4909984/.

Rutledge RB, Moutoussis M, Smittenaar P, Zeidman P, Taylor T, Hrynkiewicz L, Lam J, Skandali N, Siegel JZ, Ousdal OT, Prabhu G, Dayan P, Fonagy P, Dolan RJ (2017) Association of neural and emotional impacts of reward prediction errors with major depression. JAMA Psychiatry 74:790–797.

Rutledge RB, Skandali N, Dayan P, Dolan RJ (2014) A computational and neural model of momentary subjective well-being. Proceedings of the National Academy of Sciences 111:12252–12257.

Rutledge RB, Skandali N, Dayan P, Dolan RJ (2015) Dopaminergic modulation of decision making and subjective well-being. The Journal of Neuroscience 35:9811–9822.

Smith DV, Hayden BY, Truong T-K, Song AW, Platt ML, Huettel SA (2010) Distinct Value Signals in Anterior and Posterior Ventromedial Prefrontal Cortex. J Neurosci 30:2490–2495.

Son LK, Metcalfe J (2005) Judgments of learning: Evidence for a two-stage process. Memory & Cognition 33:1116–1129.

Son LK, Sethi R (2006) Metacognitive control and optimal learning. Cognitive Science 30:759–774.

Spaniol J, Di Muro F, Ciaramelli E (2019) Differential impact of ventromedial prefrontal cortex damage on “hot” and “cold” decisions under risk. Cogn Affect Behav Neurosci 19:477–489.

Steptoe A, Deaton A, Stone AA (2015) Subjective wellbeing, health, and ageing. Lancet 385:640–648.

Sugawara SK, Tanaka S, Okazaki S, Watanabe K, Sadato N (2012) Social Rewards Enhance Offline Improvements in Motor Skill. PLOS ONE 7:e48174.

Taylor M, Creelman CD (1967) PEST: Efficient estimates on probability functions. The Journal of the Acoustical Society of America 41:782–787.

Van de Mortel TF (2008) Faking it: social desirability response bias in self-report research. Australian Journal of Advanced Nursing, The 25:40.

Vinckier F, Rigoux L, Oudiette D, Pessiglione M (2018) Neuro-computational account of how mood fluctuations arise and affect decision making. Nature communications 9:1–12.

Wilson T, Gilbert DT (2005) Affective forecasting: knowing what to want. Psychol Sci 14:131–134.

Winecoff A, Clithero JA, Carter RM, Bergman SR, Wang L, Huettel SA (2013) Ventromedial Prefrontal Cortex Encodes Emotional Value. J Neurosci 33:11032–11039.

Zald DH, Andreotti C (2010) Neuropsychological assessment of the orbital and ventromedial prefrontal cortex. Neuropsychologia 48:3377–3391.

Zald DH, Mattson DL, Pardo JV (2002) Brain activity in ventromedial prefrontal cortex correlates with individual differences in negative affect. PNAS 99:2450–2454.

